# Large language models unlock the ecology of species interactions

**DOI:** 10.64898/2026.02.06.704115

**Authors:** Heng-Xing Zou, Xiaohao Yang, Thabassum H. Hajamaideen, Olivia J. Stein, Roxanne S. Beltran, Benjamin G. Freeman, Mark Lindquist, Eliot T. Miller, Summer Mengarelli, Charlotte M. Probst, Fernanda S. Valdovinos, Derek Van Berkel, Phoebe L. Zarnetske, Brian C. Weeks, Kai Zhu

## Abstract

Species interactions shape population dynamics, geographic distributions, evolutionary trajectories, and responses to environmental change. Yet data on these interactions remain scarce across broad spatial, temporal, and taxonomic scales because they are difficult to collect in the field. One promising source of interaction data is citizen science platforms, which contain billions of biodiversity observations, often accompanied by unstructured text comments that may document interactions among organisms. Advances in large language models (LLMs) make it increasingly feasible to identify, extract, and categorize biotic interactions from these unstructured data at scale. Here, we present an LLM workflow that collects species interaction observations from multilingual citizen science comments. Using two case studies—bird-bird and plant-pollinator interactions— we show that LLMs can rapidly extract interaction types and participating species with high accuracy. These data can greatly expand the spatial, temporal, and taxonomic coverage and resolution of species interactions data, enable new tests of long-standing ecological questions, and improve our ability to track ecological changes. With appropriate validation, expert review, and attention to data privacy for both users and sensitive species, this approach opens new opportunities to characterize, forecast, and conserve biodiversity under global change.

## Species interactions: a “new frontier” of ecological data

Biotic interactions between species are fundamental to characterizing and predicting ecosystem resilience (1), community dynamics (2), biodiversity patterns (3), ecosystem functioning (4), and evolution (5). Although large-scale data on species abundance and distribution are increasingly available (6), knowledge of species interactions remains limited across many ecosystems worldwide (7, 8). Widespread changes in geographic range and phenological shifts under rapid global change have profound consequences for species, communities, and interactions. For instance, range expansion may introduce new species into communities, creating novel interactions, whereas range contraction can lead to the local extirpation and the loss of existing interactions (9–11). Despite ample case studies, the novel ecological interactions produced by “species on the move” (i.e., shifts in spatial range and phenology) (12) have not been examined on broad spatial, temporal, and taxonomic scales. Understanding the consequences of these large-scale changes in species interactions is a major frontier in ecology, but progress is constrained by the scarcity of detailed behavioral and interaction data (13).

Such a critical biodiversity data gap persists because species interaction data remain scarce across broad spatial and temporal scales, largely due to the logistical challenges of field-based data collection (7, 14). Traditional efforts rely heavily on expert observations or experiments on a local scale, which are time-consuming and costly, making it challenging to document the full range of interactions that a focal species may experience, leading to spatial, temporal, and taxonomic constraints on resulting datasets (7, 8, 15). Because species interactions are often highly context-dependent (16, 17), small-scale datasets provide only a limited view of ecological communities, constraining our ability to generalize dynamics and predict how interactions may shift under global change.

Here, we propose that citizen science platforms, such as iNaturalist (inaturalist.org) and eBird (ebird.org) (18), which collectively contain huge amounts of biodiversity observations, can help alleviate the longstanding scarcity of species interaction data. In addition to manual screening of observation data, we highlight the power of automated data collection enabled by advances in large language models (LLMs) through two case studies on bird-bird and plant-pollinator interactions. Finally, we discuss the great potential of this vastly expanded data source to answer long-standing ecological questions and inform biodiversity conservation.

### Citizen science as a rich source of species interaction data

Citizen science engages members of the public in scientific data collection, generating vast quantities of ecological observations that increasingly support biodiversity research and conservation science (19, 20). This practice has also been termed “participatory science” (21). In this paper, citizen science specifically refers to volunteer-generated biodiversity data, which is different from community science that traditionally refers to grassroots initiatives aiming to benefit local communities (see (22, 23) for a more in-depth discussion of the terminology). Observations contributed by participants from around the world have greatly advanced our understanding of biodiversity from species to community levels across diverse groups of taxa and in both populous and remote regions (24–27). At finer scales, data from popular platforms have generated new insights into plant-frugivore (28), plant-pollinator (28–30), and predator-prey interactions (31), and even revealed previously undocumented interactions (32, 33). Beyond individual case studies, the extensive spatial and temporal coverage of citizen science data has enabled comparative analyses of species interactions across analogous ecosystems (e.g., urban ecosystems) (34, 35) and broad environmental gradients (e.g., latitudinal) (36). These data complement observations by trained scientists by expanding the spatial and taxonomic scope of documented interactions and by revealing previously unrecorded interactions, especially in highly biodiverse but often data-deficient areas (e.g., the Neotropics) (28, 30, 31, 33, 35).

Although citizen science has generated valuable data on species interactions, most platforms were originally designed to record the presence and abundance of a single taxon rather than interactions among species. The study of interactions in these datasets has largely relied on the incidental recording of observers either through optional text posts (“comments”), or through uploaded photos or other media types (19). Previous studies used these unstructured data to identify interactions, but the large volume of records has generally limited analyses to manual examination of specific interactions (e.g., plant-pollinator) within relatively small areas (e.g., in one or several metropolitan areas)(28, 29, 31). Efforts are being made to enable the structured submission of species interaction data on some platforms. For example, iNaturalist now provides functionality to explicitly record interactions (37), such as pollination, predation, and parasitism, among others. These records are automatically imported into the Global Biotic Interactions Database (GloBI) (38), where they are made readily accessible for ecological research. Project FeederWatch, a platform designed to record birds in backyard feeders (39), includes standardized submission of bird-bird interactions (displacement or predation) (40–42). Nevertheless, explicit recordings of species interactions remain uncommon across platforms, as most prioritize simplified reporting protocols that encourage participation and the collection of focal biodiversity data (19).

Despite their promise, current methods of collecting species interaction data from citizen science platforms have yet to realize the full potential of the unprecedented spatial and temporal coverage, resolution, and sheer volume of observations contributed by participants worldwide. For example, we estimate that about 3-5% of eBird observations contain comments, and 3-5% of comments contain species interactions. With over two billion observations on eBird worldwide (43), these rough estimates suggest there are ∼5,000,000 comments that contain species interaction data. Even though information on species interactions is relatively uncommon, because of the sheer volume of comments, they are a data source with great promise.

### Automating the collection of species interactions with LLMs

Collecting and analyzing species interaction data from citizen science platforms has become substantially easier due to increasing data accessibility and recent advances in natural language processing, particularly LLMs. Previous bottlenecks in obtaining citizen science observations have been alleviated through centralized biodiversity data infrastructures. For example, the Global Biodiversity Information Facility (GBIF) (44) provides access to observations from multiple platforms, including iNaturalist and observation.org. Researchers can query records by species, location, and time period, and filter observations containing comments. These records can be further filtered and processed using LLMs. Depending on the application, LLMs can be trained on manually labeled datasets (e.g., comments that contain species interactions or certain types of interactions) or used in a zero-shot or few-shot framework to identify and classify interactions from unstructured text. Here, we illustrate one potential basic training workflow (Figure 1) with case studies of bird-bird interactions and plant-pollinator interactions. LLM prompts for each case study are included in the Supplementary Material.

**Figure 1.**
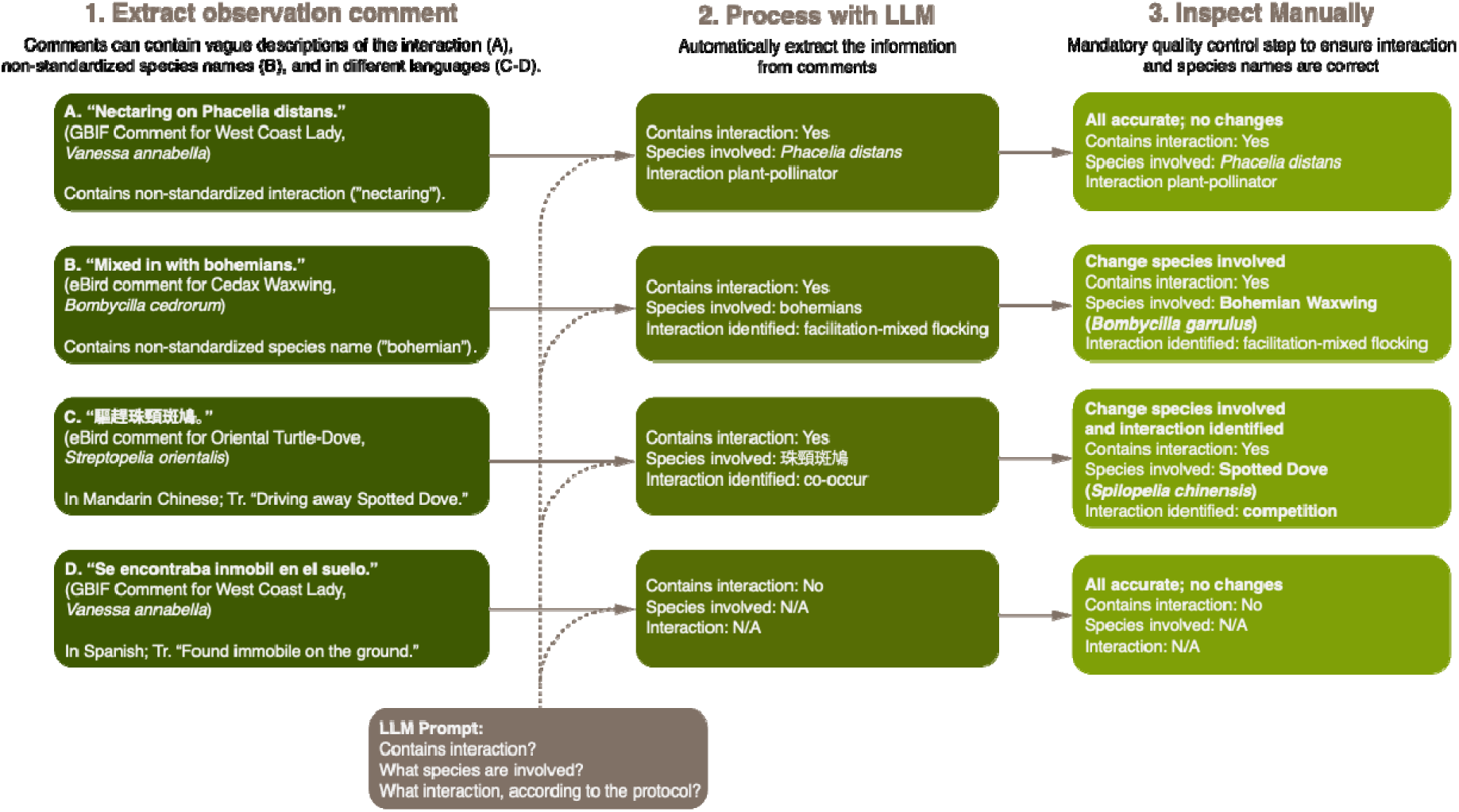
Basic workflow of extracting species interactions using LLMs. Step 1: Examples of comments from eBird and GBIF. They represent several characteristics of the comments: vague descriptions of interactions, non-standardized species names, different information density (may or may not contain interactions), and multilingual. Step 2: LLMs can extract the three key types of information in comments of multiple languages: whether a comment contains an interaction, what species were involved in the interaction, and what type of interaction was recorded. To ensure the efficiency of extraction, the non-standard species names are reported as is. Step 3: Processed comments are then manually inspected to standardize species names according to the specific taxon and to correct any misidentified interaction categories. This step is essential for quality control.

### Case study 1: eBird comments

To illustrate the potential of LLMs for extracting species interactions from citizen science platforms, we first fine-tuned a GPT-4o model (version number ft:gpt-4o-2024-08-06; see Supplementary Material for model parameters including bird-bird interaction definitions from the AvianMetaNetwork) (45) on 544 randomly selected eBird comments from the state of Texas (retrieved Aug 2023). The training dataset consisted of an equal number of comments containing species interaction information and comments without interaction information. Within the test set, the fine-tuned model correctly identified 76% of all comments containing interactions and 96% of comments that did not. We then applied the model to eBird observations from Michigan collected between January 2019 and January 2024. Of the 18,150,523 records, 632,169 (3.5%) contained comments, and the model identified 29,345 (4.6%) that contain species interactions. The cross-state application also provided a preliminary test of the model’s ability to generalize to comments originating from a different geographic region than the training data.

To further test the LLM’s accuracy in categorizing species interactions, a single annotator (O.J.S.) manually reviewed and categorized 485 comments randomly selected from those identified as containing interactions in the Michigan dataset. Using the AvianMetaNetwork protocol (45), the annotator categorized interaction types and the species involved, completing the task in approximately eight hours. We then imported the comments into Claude Sonnet 4 (version number claude-sonnet-4-20250514) with a prompt containing the categorization protocol and matched the output to the manually labeled interactions. The LLM was not trained with prior data (zero-shot) and correctly categorized 395 out of 485 (81.44%, Wilson’s interval [77.63%, 84.75%]) interactions in less than five minutes. Species were identified by the LLM as the name used in the original comments, which, in many cases, was not the standard common name. For example, the comment “Mixed with bohemians” refers to the species Bohemian Waxwing (Bombycilla garrulus) as “bohemian” (Figure 1, example comment A). After manually correcting these species names to the 2025 version of eBird taxonomy (46), the LLM correctly extracted species identity from 468 (96.49%, [94.33%, 97.88%]) comments, up from 80% before the manual correction, including comments with multispecies interactions (i.e., the focal species interacted with more than one other species).

We then constructed and compared matrices from manually labeled and LLM-extracted interactions and calculated their differences using a series of dissimilarity metrics (47): β_s_ defines the dissimilarity in species compositions; β_OS_ defines the dissimilarity of interactions between species shared by the two matrices; β_WN_ defines the overall dissimilarity of interactions; β_ST_ defines the dissimilarity of interactions due to turnover of species between the two matrices. Smaller values indicate higher similarity. For some specific interaction types, such as mobbing (aggression towards potential predators) and predation, the LLM correctly identified all interactions from the comments (Table 1). For facilitation-feeding (multiple species feeding together) (45), the labeled and extracted matrices based on the presence of species and interactions are identical, but the LLM failed to extract one interaction (Figure 2). For two common types of species associations, mixed-species flocking (multiple species benefit from being in a group) and co-occurring (spatial and temporal co-occurrence), the manually labeled and LLM-extracted matrices show high similarity (Table 1). Some common interactions are more robust to data extraction because they have been recorded in many comments, while others are more incidental but may indicate rare, understudied cases.

**Figure 2.**
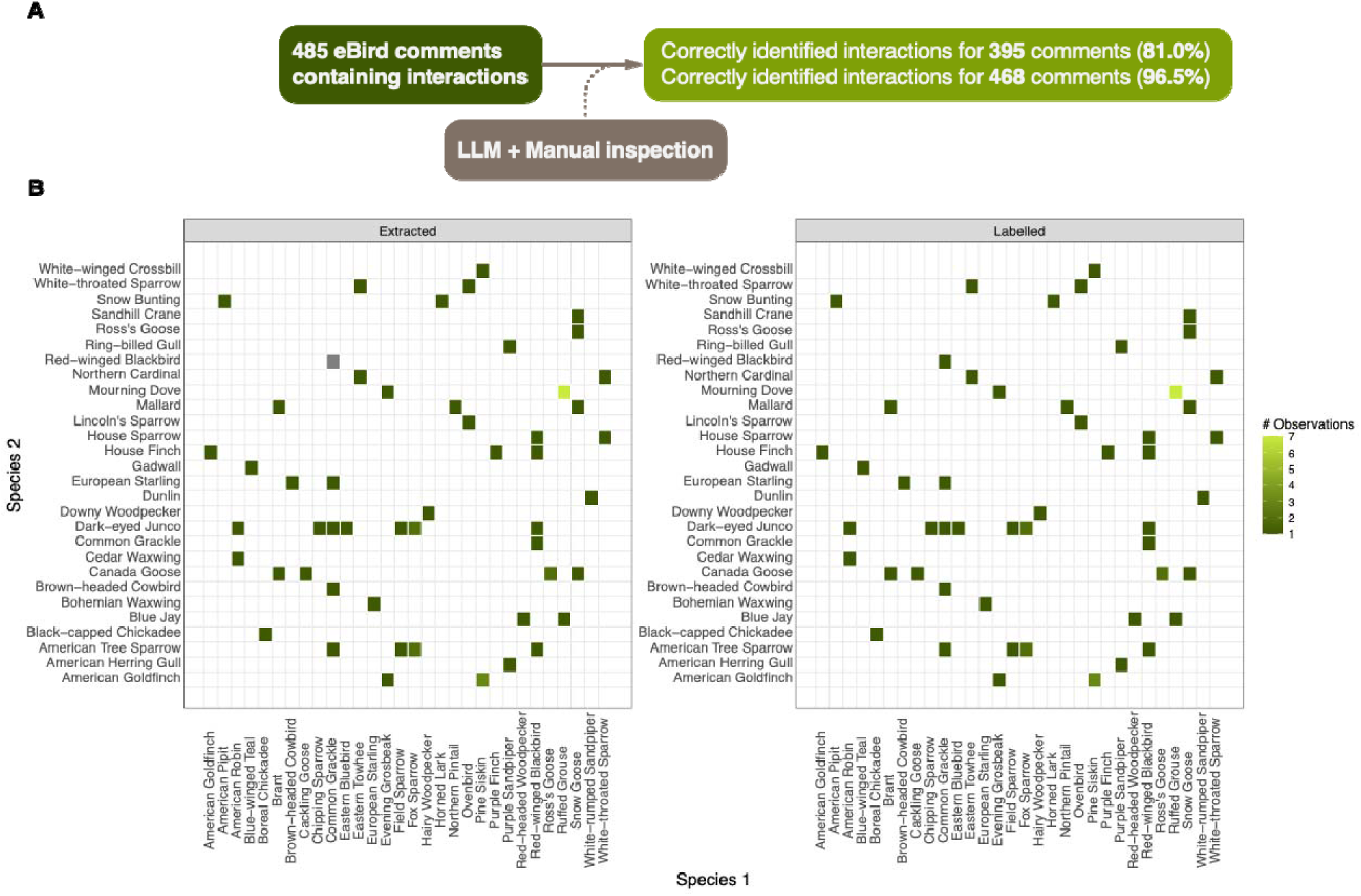
Interaction matrices of bird species feeding together in Michigan, winter 2023 and 2024. Panel A shows the analysis pipeline. Panel B left: the interaction matrix extracted from the LLM; panel B right: the interaction matrix from comments that were manually categorized according to the same protocol. The two matrices are extremely similar, and the LLM failed to extract only one interaction from a comment with Common Grackle (Quiscalus quiscula) as the focal species and mentioned Red-winged Blackbird (Agelaius phoeniceus) as a participating species (gray square on the left panel). However, this difference did not reflect on the structure of interaction matrices based only on species identity and interaction presence (i.e., non-directional, unweighted matrices; see Table I for quantitative comparison) because the interaction between the two species was identified in other comments. Overall, this result indicates that the LLM can accurately extract interactions between birds from eBird comments.

**Table 1.** Dissimilarities between interaction matrices constructed from manually labeled interactions vs. LLM-extracted interactions, following (45).

| Interaction type | $\beta_s$ | $\beta_{os}$ | $\beta_{WN}$ | $\beta_{ST}$ | # interactions in manually labeled matrices | # interactions in LLM-extracted matrices |
| --- | --- | --- | --- | --- | --- | --- |
| facilitation-feeding | 0 | 0 | 0 | 0 | 63 | 62 |
| facilitation-mixed flocking | 0 | 0.00357 | 0.00357 | 0 | 283 | 279 |
| predation | 0 | 0 | 0 | 0 | 13 | 13 |
| co-occur | 0.0253 | 0 | 0.0148 | 0.0148 | 103 | 100 |
| mobbing | 0 | 0 | 0 | 0 | 21 | 21 |
| amensalism | 0 | 0 | 0 | 0 | 8 | 8 |
| competition | 0 | 0 | 0 | 0 | 3 | 3 |
| aggression-potential competition | 0 | 0 | 0 | 0 | 3 | 3 |
| aggression | 0 | 0 | 0 | 0 | 6 | 6 |

In this case study, we used two LLMs (GPT 4o and Claude Sonnet 4) because the fine-tuning feature, only available for GPT, could help with identifying the small proportion of comments that contain interactions. In later use cases, including case study 2 reported below, we found that Claude Sonnet 4 could achieve comparable performance by analyzing comments directly obtained from eBird without fine-tuning. This indicates that cutting-edge LLMs may be sufficiently capable of identifying species interactions without prior training (zero-shot).

To test the performance of LLMs in relatively data-deficient, non-English-speaking areas, we also analyzed species interactions from 10,000 comments in Taipei, a sub-tropical city with relatively few eBird comments (62,356 comments) and a lower proportion of comments containing species interactions (1.43% in the sampled subset), with most of the comments in Chinese (see Figure 1, example comment C; see also Supplementary Material for detailed methods). The LLM successfully identified all but four comments not containing interactions (false positive rate 0.04%). Among the 143 comments with interactions, 20 were identified as no interactions (a false negative rate of 14.00%), and an additional 35 comments were identified as different interactions, leading to an overall accuracy of 61.54% (88/143, Wilson’s interval [53.00%, 69.44%]). Examining the errors, the model appeared to perform poorly in identifying bird species in Chinese, and the lack of bird species in comments led to the model concluding no interactions were recorded.

### Case study 2: Comments regarding a single pollinator from GBIF

To illustrate how LLMs can help collect plant-pollinator interactions from citizen science platforms, we downloaded all human observations of a single pollinator, the West Coast Lady (Vanessa annabella), from the Global Biodiversity Information Facility (GBIF) on Apr 25, 2025. Out of the 8,999 observations, 996 (11.07%) contained comments in both English (934 comments) and Spanish (62 comments; see Figure 1, example comment D). First, we tested the accuracy of LLMs in categorizing interactions. We manually annotated 400 comments, categorizing all interactions involving the focal species as pollination, herbivory, co-occurrence, and no interactions (Supplementary Material). Using the same protocol, we constructed a prompt for Claude Sonnet 4, then asked the LLM to categorize the interactions and extract the other species involved in the interaction. The LLM was not trained on prior data. Using the manually annotated data as validation data, we found that the LLM correctly identified 368 out of 400 (92.00%, Wilson’s interval [88.78%, 94.38%]) of all interaction types, and 396 out of 400 (99.00%, [97.28%, 99.68%]) of all species involved after correcting for non-standard species names (common names, spelling errors, etc.). This subset contained 18 comments in Spanish, and we found no sign of difference in accuracy for identifying interaction types or species involved (Chi-square test, both df = 1, p > 0.80), although the sample sizes were too small to systematically test for the potential difference among languages.

Then, we used the LLM to extract and categorize interactions from all 996 comments, yielding 299 (30.02%) comments that contained interactions, 100 of which contained observations of pollination. To compare the completeness of this plant-pollinator dataset, we downloaded all interactions involving Vanessa annabella visiting flowers from the Global Biotic Interactions Database (GloBI) (38); the GloBI subset included interactions between V. annabella and 138 unique plant species. We converted the common names of those plants in comments to their scientific names (Figure 3), then aligned them to the GBIF backbone taxonomy (48) used by GloBI. We found 46 species matching the GloBI species list, 36 taxa that matched the GloBI species list at a genus level (i.e., different species in the same genus, or species only identified to genus; hereafter “genus-match”), and 20 taxa that did not match any taxa in the GloBI species list (hereafter “no-match”; Figure 3).

**Figure 3.**
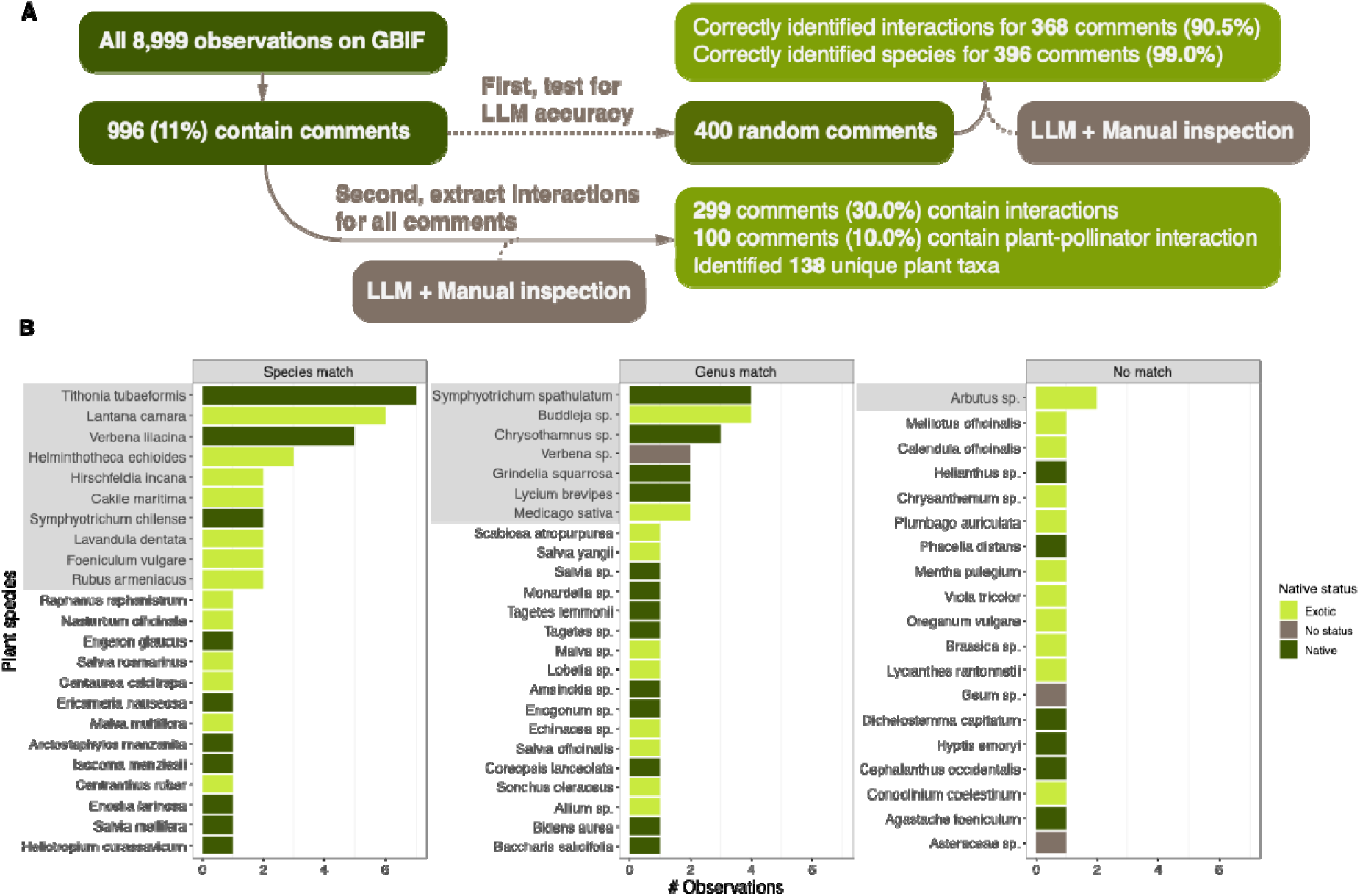
Plant-pollinator interactions of the West Coast Lady butterfly (Vanessa annabella) collected from comments on citizen science platforms. Panel A shows the analysis pipeline. In panel B, the x-axis shows the number of comments observing the interaction. The three columns indicate whether the species also appears in the GloBI species list for the plants visited by Vanessa annabella: “species match” means that the plant species in comments exactly matches a species in GloBI, “genus match” means that the plant species matches to the genus level (different species in the same genus, or species only identified to genus), and “no match” means that the plant species is not in the GloBI species list. Species are colored by their native status in the butterfly’s range (western United States and Mexico); “no status” indicate the identified taxon is too broad for determining native status. Species shaded in gray appear in more than one observation, indicating their potential ecological importance.

Although the frequency of reports in this dataset does not necessarily reflect the actual frequency of visitation by the butterfly, the presence/absence of specific species or groups can still reveal important ecological insights. From these results we made three ecologically important observations. First, among the top 10 species with the greatest number of observations, only three are native to the range of the butterfly (western United States and Mexico). Exotic species are even more prevalent among no-matches, with only six native taxa on the list. The abundance of exotic species on this list corresponds to the global pattern of introduced species integration into plant-pollinator networks (49), and the high proportion of exotics among no-matches indicates that observations on GloBI may have overlooked or underestimated introduced plant species used by the butterfly (i.e., “novel” plant-pollinator interactions that can potentially rewire the interaction networks). Second, we observed genus-matches of common ornamental plants, such as butterfly bush (Buddleja sp.) and verbena (Verbena sp.), likely because observers are often unable to determine species-level scientific names of these ornamental cultivars. These observations highlight the spatial bias of GBIF data toward residential areas in urban and suburban landscapes. Third, the genus-match with the most observations, western mountain aster (Symphylotrichum spathulatum), is a native species with a congener (Pacific aster [Symphyotrichum chilense]) that is among the top 10 of the exact matches. This indicates that data from citizen science observations may have recorded an important native plant associated with this butterfly absent in GloBI. Overall, this case study highlights that GloBI data, compiled from a wide range of studies and sources and often considered a more complete data source of interactions, can still be improved by the additional data collected from GBIF comments. Furthermore, the gap between GloBI and data extracted from GBIF comments may indicate important gaps in ecological knowledge.

### Advantages, limitations, and recommendations

In both case studies, LLMs were remarkably fast, accurate, and had high batch-processing ability (processing 500 comments in less than five minutes). Significantly, a further advantage of LLMs is their ability to handle comments in different languages in both of our case studies (Figure 1). Because citizen science platforms are widely used around the world, the ability to automatically process multiple languages combats traditional biases against non-English data sources and opens broader possibilities for the research of species interactions globally (50). Compared to other AI-enabled methods, such as extracting species interactions from computer vision (51), analyzing text using LLMs has lower computational costs, requires less domain knowledge in AI, and draws information from a large amount of data—writing textual comments is less cumbersome than taking and uploading photos to some citizen science platforms. The latter is especially relevant for eBird since research-grade bird photos often require high-end cameras inaccessible to many observers. While LLMs can also extract species interactions from published scientific papers (15), compared to the wide spatial, temporal, and taxonomic coverage from millions of users around the world, scientific literature is more subject to the same data shortfall of previous datasets on species interactions (7, 8) and may not fully represent the spatial and temporal turnover of interactions. Together, the LLM-based analysis of citizen science comments has the unique potential to revolutionize the collection of interaction data, greatly advancing research on topics from classic ecological questions to emergent issues under global change.

Through the case studies, we identify three major limitations of LLMs. First, results from the model may not be consistent among different runs (e.g., one comment could be classified as having different types of interactions by the same model) (52). This can be improved by training the model with manually labeled data to improve accuracy, running the model several times, crafting unambiguous prompts with direct instructions, and, most importantly, manually inspecting the model output. Second, LLMs sometimes cannot categorize the interactions or infer standard common names of participating species mentioned in comments, which often contain vague language to describe interactions or when referring to species (Figure 1). This can be improved by providing more specific prompts that are supplied to the model (e.g., listing keywords that may appear in comments that describe certain interactions or specific examples; Supplementary Material), manual inspection, or cross-checking with videos or photos associated with the same observation, either manually or by using deep learning models specializing in computer vision (53). Third, our case studies revealed considerable differences in the accuracy of LLMs among comments in English and Chinese, but not among English and Spanish. This difference in performance among human languages is especially pertinent because data-deficient regions for species interactions often also use languages that are not well-parsed by major LLMs (50, 54). In our case study on eBird comments from Taipei, the low performance might be due to a lack of training text on bird names in Chinese. This gap can be partially mitigated by fine-tuning the model in collaboration with local experts who use the language, which can also in part address the global imbalance in biodiversity data and ecological research (55).

We also flag two common issues of LLMs that are important to consider in other use cases. First, LLMs can generate content that is absent in the input, a common problem called hallucination (56). We did not find any obvious cases of hallucination in our case studies, partly because we gave detailed instructions in our prompt to extract information from short input (comments) into a short, standardized format (either “Yes” or “No” for identifying species interactions, or a set of choices for categorizing species interactions). This rigorously formatted data extraction is less prone to hallucination, which usually arises from longer content generation. Second, output from LLMs can be highly sensitive to the wording of the prompt (57). In our case studies, we did not find large differences in results from different iterations of the prompt during testing, partly due to the specific categorization protocols that include common keywords indicating certain types of interactions and the standardized output format.

In our case studies, we used closed-source models such as GPT 4o and Claude Sonnet 4. Compared to open-source models, these models may be subject to changes without notice, limiting the reproducibility of analyses. The gold standard of scientific research is reproducibility, and to achieve this, we encourage researchers to use open-source LLMs configured locally (offline), allowing for better customization, higher consistency, and lower resource requirements that reduce environmental impacts. We tested the performance of two open-source models, LlaMA 3 8B and Mistral 7B v0.2, and found that using the same test dataset (eBird comments used in case study 1), the accuracy of categorizing species interactions was much lower even after fine-tuning (55.3% and 42.8%; see Supplementary Material for details). Although open-source models produced less accurate results at the time of this test (May 2025), with recent technological advances, we encourage researchers to explore open-source options whenever possible.

In addition to general guidelines for using AI tools in ecological research (14, 58, 59), we provide the following recommendations for obtaining, extracting, and curating species interaction data from citizen science comments. Citizen science data have spatial, temporal, and taxonomic biases and are inherently subject to observation errors (60–62). One specific limitation is the bias toward easily observable, “charismatic” taxa in citizen science datasets, such as the birds and butterflies covered in our case studies. Measures should be taken to address the widespread detection biases and errors of species interactions in citizen science data (e.g., higher detection for more common and more conspicuous species and their interactions, or misidentification of species interactions) using statistical methods that explicitly correct for these biases (63, 64). When obtaining the data, we recommend setting clear research objectives and questions limited to specific species, interactions, or spatial and temporal extents. Clear objectives will help design specific prompts that improve accuracy and limit the amount of data needed to be processed by LLMs and manually, which ensures the benefit of large-scale data analysis with a reasonable computational cost and workload. When extracting data, we recommend clear, detailed prompts with rigorous biological definitions of targeted interactions and short, structured output to reduce hallucination and variability in LLMs. This step is key to the performance of data extraction and requires customization and ample testing, ideally with a manually labeled dataset as illustrated in both case studies above. When curating and publishing data, researchers should adhere to the terms and conditions of specific citizen science platforms and implement measures to protect user privacy and sensitive species. For instance, eBird and iNaturalist terms do not explicitly prohibit the reuse of materials for AI tools (65, 66) and will hide coordinates of sensitive species unless requested (67, 68). Other measures of privacy protection, such as obscuring exact coordinates by a small buffer and redacting names of observers and specific locations, should be implemented when appropriate. Finally, we recommend that all output should be reviewed by experts with domain knowledge to ensure the accuracy of extracted data. While part of the advantage of this approach is the possibility of revealing previously unrecorded interactions, as showcased by case study 2, the possibility of these interactions should be checked by domain experts before further analysis.

Outlook: What can large-scale interaction data tell us?

Here, we outline several research themes that can be greatly accelerated by extracting large-scale data on species interactions from citizen science data using LLMs.

### Biogeographical patterns of species interactions

Many classic themes of biogeography center on species interactions, such as the role of biotic interactions in determining species ranges (69), and latitudinal and elevational gradients of interaction strengths (70). Past studies that have characterized these macroecological patterns relied on datasets collected from a few representative sites and focused on a specific interaction. For example, latitudinal patterns of seed predation and insect predation can be derived from spatially replicated distributed experiments, but they often lack taxonomic information of predators (70, 71). Other studies have inferred the role of species interactions, such as competition, in shaping geographical ranges across latitudinal or elevational ranges using observed and modeled species distributions (69, 72). While these approaches have generated valuable insights into the role of species interactions in shaping biogeographic patterns, they lack spatial and taxonomic coverage, diversity of interaction types, identity of specific species involved, or direct evidence of species interactions. After correcting for spatiotemporal biases, data collected from citizen science platforms could address these shortfalls by providing high-resolution, human-observed interaction data across large spatial scales and over time. In addition, these data can also advance our understanding of how interactions are likely to shift over time with global change (73), for example, with geographical range shifts of participating species (74), or along urban-rural gradients (75). Together, these large-scale species interaction data pave the first steps into building a “biogeography of species interactions” (76).

### Structure and turnover of interaction networks

Interaction networks describe how species in diverse communities are connected via different types of interactions, and these networks are essential to understanding the energy flow and stability of communities (77). Due to logistical constraints, interaction networks constructed using expert observations can often only focus on spatial coverage, temporal coverage, or the resolution of species interactions. For example, observations of plant-pollinator interactions over time provide detailed information on their temporal turnover (78) but are limited in both taxonomy (e.g., only including flowering plants and their pollinators) and space (data only taken from a few dedicated sites). On the other hand, a few highly resolved interaction networks aim for complete coverage of focal taxa and their interactions in single communities (77), but they lack coverage in both space and time. As the strengths and directions of species interactions can change across space (73), time (79–81), and environmental conditions (82, 83), these fragmented networks can only provide incomplete pictures of interaction networks in nature (84). With broader spatial, temporal, and taxonomic coverage of species interaction data, it would be possible to expand both the scope and resolution of interaction networks greatly. This expansion would allow for analyses of the turnover of interaction networks (78, 85, 86), evaluating the effect of time, space, and global change factors in “rewiring” biotic interactions and shaping ecological communities.

### Conservation and discovery of biodiversity

Natural history observations spur discoveries in ecology and can promote more effective conservation. With increasing coverage across space, time, and taxonomy, citizen science platforms can expand natural system monitoring programs (19). For example, there is little overlap between plant-hummingbird interactions collected by experts and those recorded on citizen science platforms in Brazil (29), indicating the latter’s great potential in revealing previously unrecorded interactions even among well-studied taxa. Examinations of citizen science data have also led to the discovery of cockroaches as pollinators in Spain (32) and the previously undocumented nectar-robbing behavior of a hummingbird in Bolivia (33). These discoveries can facilitate the description and conservation of biodiversity and are especially important in diverse but data-deficient areas such as the tropics. Data from citizen science platforms can also assist conservation practices. For example, some comments may report information about banded individuals or individuals with tracking devices, which is useful for tracking the focal animal’s location, condition, and biotic interactions (87). Furthermore, novel interactions arise when invasive alien species expand into new habitats. Citizen science has long been useful for monitoring the spread and consequences of invasive alien species (88, 89), and these numerous observations likely contain documentation of their interactions with native species that can be used to inform invasive alien species management.

### Support for other existing tools and data

Recent progress in statistical modeling and non-generative machine learning has provided some solutions for ecologists to overcome species interaction data limitations. Large-scale interaction data collected from citizen science platforms can further expand and support these tools. Statistical models such as joint species distribution models enable the extraction of covariances, or even interaction strengths, from time series data of species abundances across space (82, 90). However, these statistical associations between species abundances could arise from many mechanisms other than interactions and thus do not reflect true underlying interactions (91, 92). Data on recorded species interactions can be used to inform such model-fitting efforts: if two species are known to compete from data records, then the negative associations between their abundance or distribution may be a consequence of this negative interaction. Non-generative machine learning models, such as random forest and deep learning models, can predict species interactions from existing interaction networks and other data that may influence the potential of species interactions: for example, species with certain matching functional traits are more likely to interact (8, 93, 94). However, these machine learning models still require a large amount of training data and are therefore limited by the low availability of species interaction datasets. Large-scale interaction data from citizen science platforms can provide more data for the training process, improving the accuracy of such predictions (8).

### Outstanding questions

Here, we provide several outstanding questions that are relevant to both ecologists and data scientists, from broad to focused. We hope these questions can spur new and exciting research using this novel approach.

1. How can data collected from comments by LLMs address the rapid “rewiring” of species interaction networks in regions experiencing profound environmental changes?
2. How do the frequency, diversity, and structure of species interactions vary across urbanization gradients, and can citizen science platforms uncover consistent patterns across cities worldwide?
3. Can species interaction observations extracted from citizen science comments improve our understanding of impacts from disturbances, such as wildfire, drought, habitat fragmentation, or species invasions?
4. Which species groups, geographic regions, habitat types, and interaction types have less available data and thus have the potential to benefit greatly from the large amount of data collected by LLMs?
5. How can the widespread sampling bias in citizen science (spatial, temporal, geographic, taxonomic) be addressed more effectively when collecting interaction data?
6. To what extent do introduced species integrate into existing ecological networks, and are citizen science comments disproportionately capturing novel interactions involving invasive or ornamental species?
7. How can data extracted from comments help understand the interaction between humans and wildlife, both in terms of human-wildlife interaction and conflict, and human perception of nature?
8. What types of species interactions are most easily identifiable using LLMs, what types of interactions require much more nuance, and how do we address this difference in detectability among interactions?
9. How can LLMs be integrated into other advanced AI tools on topics of ecology, such as computer vision models and deep learning models that can analyze photos and videos?
10. What other fields and topics in ecology, behavior, and conservation can be addressed by the increasingly available species interaction data?

## Conclusion

Around the world, tens of thousands of observations are uploaded to citizen science platforms every hour. Data contributed to these platforms have greatly increased our understanding of species abundances and distributions. The application of LLMs to identify and analyze species interactions from submission comments has the potential to further unlock the power of citizen science data in describing and understanding species interactions, a key component in describing and predicting biodiversity. With high spatiotemporal resolution and coverage, the immense amount of data offers a great opportunity for research on classic and emerging ecological questions, ample support for existing and emerging tools of ecological analyses, and a source to inform conservation practices under a rapidly changing global environment. We believe that responsibly leveraging LLMs to collect species interaction data is a key example of how advanced AI can be used to accelerate the fields of ecology and global change biology.

## Acknowledgements

We thank Daijiang Li, the editor, and two anonymous reviewers for feedback on the manuscript. Funding was provided by the David and Lucile Packard Foundation to B.C.W. and the Michigan Institute for Data and AI in Society to H.-X.Z. H.-X.Z. was also supported by a postdoctoral fellowship from the Institute for Global Change Biology at the University of Michigan. O.J.S. was supported by the Undergraduate Research Opportunity Program at the University of Michigan. R.S.B. was supported by the David and Lucile Packard Foundation and the Alfred P. Sloan Foundation. B.G.F was supported by the David and Lucile Packard Foundation. F.S.V. was supported by NSF DEB-2129757 and DEB-2224915. P.L.Z. was supported by the Ecology, Evolution, and Behavior Program at Michigan State University. K.Z. was supported by NSF 230619 and USDA McIntire-Stennis Capacity Grant 25-PAF01509. This research was supported in part through computational resources and services provided by the Advanced Research Computing at the University of Michigan, Ann Arbor.

## Author Contributions

H.-X.Z., B.C.W., and K.Z. conceived the project. C.M.P. and S.M. also contributed to the early conception of the idea. X.Y. and T.H.H. trained and ran the large language models. O.J.S. manually scored the eBird comments used in the study. P.L.Z. provided the original protocol for categorizing bird interactions. H.-X.Z. wrote the first draft, and all other authors contributed substantially to the writing, case studies, and visualization.

## Data Availability

Original data for case studies were obtained fromeBird (ebird.org) and Global Biodiversity Information Facility (GBIF; gbif.org). Data processed from the large language model and code used for the analyses are available at github.com/hengxingzou/Zouetal2026bioRXiv and will be permanently archived upon acceptance. Prompts containing categorization protocols for species interactions are included in the Supplementary Material.

## Notes

### Competing Interest Statement

The authors have declared no competing interest.

### Summary of Updates

This revision added new test cases for the performance of LLM and more discussion on the best practices for using the proposed pipeline for data extraction.

## Reference

1. S. Haberstroh, C. Werner, The role of species interactions for forest resilience to drought. Plant Biology 24, 1098–1107 (2022).

2. J. M. Levine, J. Bascompte, P. B. Adler, S. Allesina, Beyond pairwise mechanisms of species coexistence in complex communities. Nature 546, 56–64 (2017).

3. C. Ratzke, J. Barrere, J. Gore, Strength of species interactions determines biodiversity and stability in microbial communities. Nat Ecol Evol 4, 376–383 (2020).

4. A. Gonzalez, et al., Scaling-up biodiversity-ecosystem functioning research. Ecology Letters 23, 757–776 (2020).

5. M. G. Weber, C. E. Wagner, R. J. Best, L. J. Harmon, B. Matthews, Evolution in a Community Context: On Integrating Ecological Interactions and Macroevolution. Trends in Ecology & Evolution 32, 291–304 (2017).

6. M. Dornelas, et al., BioTIME: A database of biodiversity time series for the Anthropocene. Global Ecology and Biogeography 27, 760–786 (2018).

7. J. Hortal, et al., Seven Shortfalls that Beset Large-Scale Knowledge of Biodiversity. Annual Review of Ecology, Evolution, and Systematics 46, 523–549 (2015).

8. T. Strydom, et al., A roadmap towards predicting species interaction networks (across space and time). Philosophical Transactions of the Royal Society B: Biological Sciences 376, 20210063 (2021).

9. A. J. A. Stewart, et al., The role of ecological interactions in determining species ranges and range changes. Biological Journal of the Linnean Society 115, 647–663 (2015).

10. P. E. Hulme, Climate change and biological invasions: evidence, expectations, and response options. Biological Reviews 92, 1297–1313 (2017).

11. J. C. Fowler, M. L. Donald, J. L. Bronstein, T. E. X. Miller, The geographic footprint of mutualism: How mutualists influence species’ range limits. Ecological Monographs 93, e1558 (2023).

12. A. L. Fredston, et al., Reimagining species on the move across space and time. Trends in Ecology & Evolution 0 (2025).

13. P. P. A. Staniczenko, P. Sivasubramaniam, K. B. Suttle, R. G. Pearson, Linking macroecology and community ecology: refining predictions of species distributions using biotic interaction networks. Ecology Letters 20, 693–707 (2017).

14. L. J. Pollock, et al., Harnessing artificial intelligence to fill global shortfalls in biodiversity knowledge. Nat. Rev. Biodivers. 1, 166–182 (2025).

15. F. Keck, H. Broadbent, F. Altermatt, Extracting massive ecological data on state and interactions of species using large language models. [Preprint] (2025). Available at: https://www.biorxiv.org/content/10.1101/2025.01.24.634685v1 [Accessed 24 April 2025].

16. S. A. Chamberlain, J. L. Bronstein, J. A. Rudgers, How context dependent are species interactions? Ecology Letters 17, 881–890 (2014).

17. C. Song, S. Von Ahn, R. P. Rohr, S. Saavedra, Towards a Probabilistic Understanding About the Context-Dependency of Species Interactions. Trends in Ecology and Evolution 35, 384– 396 (2020).

18. B. L. Sullivan, et al., eBird: A citizen-based bird observation network in the biological sciences. Biological Conservation 142, 2282–2292 (2009).

19. Q. Groom, et al., Species interactions: next-level citizen science. Ecography 44, 1781–1789 (2021).

20. B. M. Mason, et al., iNaturalist accelerates biodiversity research. BioScience 75, 953–965 (2025).

21. R. Mady, Why Do We Call It Participatory Science? Smithsonian Environmental Research Center (2023). Available at: https://serc.si.edu/why-do-we-call-it-participatory-science [Accessed 1 June 2026].

22. C. B. Cooper, et al., Inclusion in citizen science: The conundrum of rebranding. Science 372, 1386–1388 (2021).

23. D. E. Lin Hunter, G. J. Newman, M. M. Balgopal, What’s in a name? The paradox of citizen science and community science. Frontiers in Ecology and the Environment 21, 244–250 (2023).

24. A. G. Gaier, J. Resasco, Does adding community science observations to museum records improve distribution modeling of a rare endemic plant? Ecosphere 14, e4419 (2023).

25. J. Socolar, et al., Seasonal macro-demography of North American bird populations revealed through participatory science. Ecography 2025, e07349 (2025).

26. J. J. Chu, D. P. Gillis, S. H. Riskin, Community science reveals links between migration arrival timing advance, migration distance and wing shape. Journal of Animal Ecology 91, 1651–1665 (2022).

27. M. Fajgenblat, et al., Leveraging Massive Opportunistically Collected Datasets to Study Species Communities in Space and Time. Ecology Letters 28, e70094 (2025).

28. A. Díaz, et al., Diet and bird-plant interaction networks based on citizen science data in Lima, Peru: exotic and native species are important. Studies on Neotropical Fauna and Environment 59, 1028–1043 (2024).

29. C. Bosenbecker, et al., Contrasting nation-wide citizen science and expert collected data on hummingbird–plant interactions. Perspectives in Ecology and Conservation 21, 164–171 (2023).

30. D. De Jong, T. M. Francoy, Citizen Science and Pollinators in South America, A New Book Freely Available Online in Spanish and Portuguese. Bee World 99, 103–105 (2022).

31. E. de Souza, et al., Ophiophagy in Brazilian birds: a contribution from a collaborative platform of citizen science. Ornithol. Res. 30, 15–24 (2022).

32. Á. Pérez-Gómez, M. León-Osper, D. Pareja, J. Robla, Flower visits of cockroaches (Insecta: Blattodea) in the Iberian Peninsula: Are they neglected pollinators? Journal of Applied Entomology 147, 565–576 (2023).

33. L. Telleria, R. Calbimonte, F. Montaño-Centellas, NECTAR ROBBING BY THE RED-TAILED COMET SAPPHO SPARGANURUS: THE VALUE OF CITIZEN SCIENCE TO DOCUMENT INFREQUENT BEHAVIOR IN BIRDS. Ornitología Neotropical 35, 20–22 (2024).

34. J. W. Wei, B. P. Y.-H. Lee, L. B. Wen, Citizen Science and the Urban Ecology of Birds and Butterflies — A Systematic Review. PLOS ONE 11, e0156425 (2016).

35. O. H. Marín-Gómez, C. Rodríguez Flores, M. del C. Arizmendi, Assessing ecological interactions in urban areas using citizen science data: Insights from hummingbird–plant meta-networks in a tropical megacity. Urban Forestry & Urban Greening 74, 127658 (2022).

36. A. J. Parker, J. D. Thomson, Citizen scientists document geographic patterns in pollinator communities. Journal of Pollination Ecology 23, 90–97 (2018).

37. Observation field: Ecological interaction · iNaturalist. iNaturalist. Available at: https://www.inaturalist.org/observation_fields/1050 [Accessed 7 January 2026].

38. J. H. Poelen, J. D. Simons, C. J. Mungall, Global biotic interactions: An open infrastructure to share and analyze species-interaction datasets. Ecological Informatics 24, 148–159 (2014).

39. D. N. Bonter, E. I. Greig, Over 30 Years of Standardized Bird Counts at Supplementary Feeding Stations in North America: A Citizen Science Data Report for Project FeederWatch. Front. Ecol. Evol. 9 (2021).

40. E. T. Miller, et al., Fighting over food unites the birds of North America in a continental dominance hierarchy. Behavioral Ecology 28, 1454–1463 (2017).

41. I. Berberi, E. T. Miller, R. Dakin, The effect of sociality on competitive interactions among birds. Proceedings of the Royal Society B: Biological Sciences 290, 20221894 (2023).

42. Detailed Instructions. Project FeederWatch. Available at: https://feederwatch.org/about/detailed-instructions/ [Accessed 7 January 2026].

43. T. eBird, eBird Passes 2 Billion Bird Observations - eBird. Available at: https://ebird.org/ebird/news/ebird-passes-2-billion-bird-observations [Accessed 7 January 2026].

44. 44. What is GBIF? Available at: https://www.gbif.org/what-is-gbif [Accessed 29 July 2025].

45. P. L. Zarnetske, et al., The AvianMetaNetwork: biotic interactions among birds of the continental United States and Canada. [Preprint] (2026). Available at: https://www.biorxiv.org/content/10.64898/2026.05.11.723238v1 [Accessed 18 May 2026].

46. E. J. McTavish, et al., A complete and dynamic tree of birds. Proceedings of the National Academy of Sciences 122, e2409658122 (2025).

47. T. Poisot, E. Canard, D. Mouillot, N. Mouquet, D. Gravel, The dissimilarity of species interaction networks. Ecology Letters 15, 1353–1361 (2012).

48. GBIF Secretariat, GBIF Backbone Taxonomy. (2023). 10.15468/39omei.

49. D. B. Stouffer, A. R. Cirtwill, J. Bascompte, How exotic plants integrate into pollination networks. Journal of Ecology 102, 1442–1450 (2014).

50. T. Amano, J. P. González-Varo, W. J. Sutherland, Languages Are Still a Major Barrier to Global Science. PLOS Biology 14, e2000933 (2016).

51. M. N. Ratnayake, A. G. Dyer, A. Dorin, Tracking individual honeybees among wildflower clusters with computer vision-facilitated pollinator monitoring. PLOS ONE 16, e0239504 (2021).

52. M. Staudinger, W. Kusa, F. Piroi, A. Lipani, A. Hanbury, A Reproducibility and Generalizability Study of Large Language Models for Query Generation in Proceedings of the 2024 Annual International ACM SIGIR Conference on Research and Development in Information Retrieval in the Asia Pacific Region, SIGIR-AP 2024., (Association for Computing Machinery, 2024), pp. 186–196.

53. Z. Miao, et al., New frontiers in artificial intelligence for biodiversity research and conservation with multimodal language models. Methods in Ecology and Evolution 17, 238–256 (2026).

54. N. Unjum, S. Evert, K. Sarveswaran, R. Weerasinghe, N. Jayatilleke, LLMs for Low-Resource Languages: A Survey. [Preprint] (2025). Available at: https://papers.ssrn.com/abstract=6122026 [Accessed 12 May 2026].

55. M. Chapman, et al., Biodiversity monitoring for a just planetary future. Science 383, 34–36 (2024).

56. Z. Ji, et al., Survey of Hallucination in Natural Language Generation. ACM Comput. Surv. 55, 248:1–248:38 (2023).

57. J. Zhuo, et al., ProSA: Assessing and Understanding the Prompt Sensitivity of LLMs in Findings of the Association for Computational Linguistics: EMNLP 2024, Y. Al-Onaizan, M. Bansal, Y.-N. Chen, Eds. (Association for Computational Linguistics, 2024), pp. 1950–1976.

58. G. R. Smith, et al., Ten simple rules for using large language models in science, version 1.0. PLOS Computational Biology 20, e1011767 (2024).

59. N. Fox, E. Di Minin, N. Carter, S. Tomkins, D. Van Berkel, Balancing accessibility and security: Safeguarding citizen-sourced biodiversity data in the age of AI and open-sourced software. Ecological Informatics 92, 103443 (2025).

60. M. Kosmala, A. Wiggins, A. Swanson, B. Simmons, Assessing data quality in citizen science. Frontiers in Ecology and the Environment 14, 551–560 (2016).

61. D. Fraisl, et al., Citizen science in environmental and ecological sciences. Nat Rev Methods Primers 2, 64 (2022).

62. L. J. Backstrom, C. T. Callaghan, H. Worthington, R. A. Fuller, A. Johnston, Estimating sampling biases in citizen science datasets. Ibis 167, 73–87 (2025).

63. T. J. Bird, et al., Statistical solutions for error and bias in global citizen science datasets. Biological Conservation 173, 144–154 (2014).

64. M. A. M. de Aguiar, et al., Revealing biases in the sampling of ecological interaction networks. PeerJ 7, e7566 (2019).

65. Terms of Use · iNaturalist. iNaturalist. Available at: https://www.inaturalist.org/pages/terms [Accessed 12 May 2026].

66. How can your eBird data be used? Help Center. Available at: https://support.ebird.org/en/support/solutions/articles/48001078113-ebird-data-privacy-and-data-use [Accessed 12 May 2026].

67. How can I access the hidden coordinates of sensitive species? iNaturalist Help. Available at: https://help.inaturalist.org/en/support/solutions/articles/151000170343-how-can-i-access-the-hidden-coordinates-of-sensitive-species-[Accessed 12 May 2026].

68. Sensitive Species in eBird. Help Center. Available at: https://support.ebird.org/en/support/solutions/articles/48000803210-sensitive-species-in-ebird [Accessed 12 May 2026].

69. B. G. Freeman, M. Strimas-Mackey, E. T. Miller, Interspecific competition limits bird species’ ranges in tropical mountains. Science 377, 416–420 (2022).

70. A. L. Hargreaves, et al., Latitudinal gradients in seed predation persist in urbanized environments. Nat Ecol Evol 8, 1897–1906 (2024).

71. E. L. Zvereva, et al., Predation on Live and Artificial Insect Prey Shows Different Global Latitudinal Patterns. Global Ecology and Biogeography 33, e13899 (2024).

72. D. W. Armitage, S. E. Jones, Coexistence barriers confine the poleward range of a globally distributed plant. Ecology Letters 1–11 (2020). 10.1111/ele.13612.

73. A. L. Hargreaves, Geographic Gradients in Species Interactions: From Latitudinal Patterns to Ecological Mechanisms. Annual Review of Ecology, Evolution, and Systematics 55, 369– 393 (2024).

74. J. A. Lawlor, et al., Mechanisms, detection and impacts of species redistributions under climate change. Nat Rev Earth Environ 5, 351–368 (2024).

75. P. Moreno-García, et al., The effects of urbanization on species interactions. Nat Cities 2, 693–702 (2025).

76. W. Thuiller, et al., Navigating the integration of biotic interactions in biogeography. Journal of Biogeography 51, 550–559 (2024).

77. K. R. S. Hale, et al., A highly resolved network reveals the role of terrestrial herbivory in structuring aboveground food webs. Philosophical Transactions of the Royal Society B: Biological Sciences 379, 20230180 (2024).

78. J. M. Olesen, J. Bascompte, H. Elberling, P. Jordano, Temporal Dynamics in a Pollination Network. Ecology 89, 1573–1582 (2008).

79. L. H. Yang, Toward a more temporally explicit framework for community ecology. Ecological Research 35, 445–462 (2020).

80. H.-X. Zou, V. H. W. Rudolf, Bridging theory and experiments of priority effects. Trends in Ecology & Evolution 38, 1203–1216 (2023).

81. H. Yin, V. H. W. Rudolf, Time is of the essence: A general framework for uncovering temporal structures of communities. Ecology Letters 27, e14481 (2024).

82. J. S. Clark, C. Lane Scher, M. Swift, The emergent interactions that govern biodiversity change. Proceedings of the National Academy of Sciences of the United States of America 117, 17074–17083 (2020).

83. O. R. Liu, S. D. Gaines, Environmental context dependency in species interactions. Proceedings of the National Academy of Sciences 119, e2118539119 (2022).

84. T. Poisot, et al., Global knowledge gaps in species interaction networks data. Journal of Biogeography 48, 1552–1563 (2021).

85. J. E. Kemp, D. M. Evans, W. J. Augustyn, A. G. Ellis, Invariant antagonistic network structure despite high spatial and temporal turnover of interactions. Ecography 40, 1315–1324 (2017).

86. E. C. Fricke, J.-C. Svenning, Accelerating homogenization of the global plant–frugivore meta-network. Nature 585, 74–78 (2020).

87. J. Parham, J. Crall, C. Stewart, T. Berger-Wolf, D. I. Rubenstein, Animal population censusing at scale with citizen science and photographic identification in AAAI Spring Symposium-Technical Report, (2017).

88. J. Encarnação, M. A. Teodósio, P. Morais, Citizen Science and Biological Invasions: A Review. Front. Environ. Sci. 8 (2021).

89. A. W. Crall, et al., Assessing citizen science data quality: an invasive species case study. Conservation Letters 4, 433–442 (2011).

90. L. J. Pollock, et al., Understanding co-occurrence by modelling species simultaneously with a Joint Species Distribution Model (JSDM). Methods in Ecology and Evolution 5, 397–406 (2014).

91. F. G. Blanchet, K. Cazelles, D. Gravel, Co-occurrence is not evidence of ecological interactions. Ecology Letters 23, 1050–1063 (2020).

92. D. Zurell, L. J. Pollock, W. Thuiller, Do joint species distribution models reliably detect interspecific interactions from co-occurrence data in homogenous environments? Ecography 41, 1812–1819 (2018).

93. E. C. Fricke, et al., Collapse of terrestrial mammal food webs since the Late Pleistocene. Science 377, 1008–1011 (2022).

94. J. Llewelyn, et al., Predicting predator–prey interactions in terrestrial endotherms using random forest. Ecography 2023, e06619 (2023).

