## Supplementary Material for "Large language models unlock the ecology of species interactions"

**Case study 1: eBird comments**

The eBird case study consists of three steps: (1) fine-tuning an LLM to identify comments that contain bird interactions using a small training set, (2) applying the fine-tuned model to the focal dataset, and (3) applying an LLM to categorize the interactions in a small subset of the focal dataset and evaluating its performance. Below, we describe the methods for steps 1 and 3 in detail.

*Fine-tuning an LLM to identify comments that contain bird interactions*

In August 2024, we downloaded all eBird observations in the state of Texas, then manually selected and categorized 544 comments, including equal proportions of comments that contain vs. do not contain interactions. An example of a comment containing interaction is

“*Being mobbed by two crows*”,

and an example of a comment that does not contain interaction is

“*foraging in adjacent lot*”.

We fine-tuned a GPT-4o model using this dataset.

For model fine-tuning, our dataset preparation process ensured a balanced, transparent, and traceable workflow for subsequent model development and evaluation. In the dataset of bird interaction, we labeled the presence of interaction as “Yes” and the absence as “No”. The dataset was split into two sets for model training and evaluation. The original dataset of 544 comment samples was first shuffled to remove any ordering bias, and the samples were then divided into training (70%), validation (10%), and testing (20%)

sets. To maintain class balance, the training and testing sets were constructed by randomly sampling equal proportions of “Yes” and “No” samples from the labeled data. To finetune the GPT model via OpenAI API (https://platform.openai.com/docs/api-reference), each record was formatted into a standardized JSONL structure containing the text content and its corresponding label, guided by pre-defined instruction templates for model fine-tuning and testing (i.e., “prompt”):

I need help classifying eBird comments by whether they describe interactions between bird species or not. Please answer with "yes" if the comments describe an interaction and "no" if not.

I am only interested in interactions between different bird species, so please exclude intraspecific interactions (interacting individuals of the same species) or interactions with other organisms that are not birds (mammals, insects, fish, amphibians, plants).

I define interactions as either direct engagement (e.g., mobbing, predation, aggression, nest guarding, flocking, parasitism) or co-occurrence (when different species co-occur in the same habitat or in close physical proximity). Please note that your answer should be only "yes" or "no".

With a learning multiplier of 0.05, temperature of 0.2, and k-top of 0.8, the GPT-4o model was trained in 4 epochs. As the learning proceeded, training and validation losses demonstrated a clear downward trend and became stable at approximately 0.5, indicating effective learning of the model (Fig. S1). The close alignment between the two curves toward the end suggests minimal overfitting and good generalization performance.

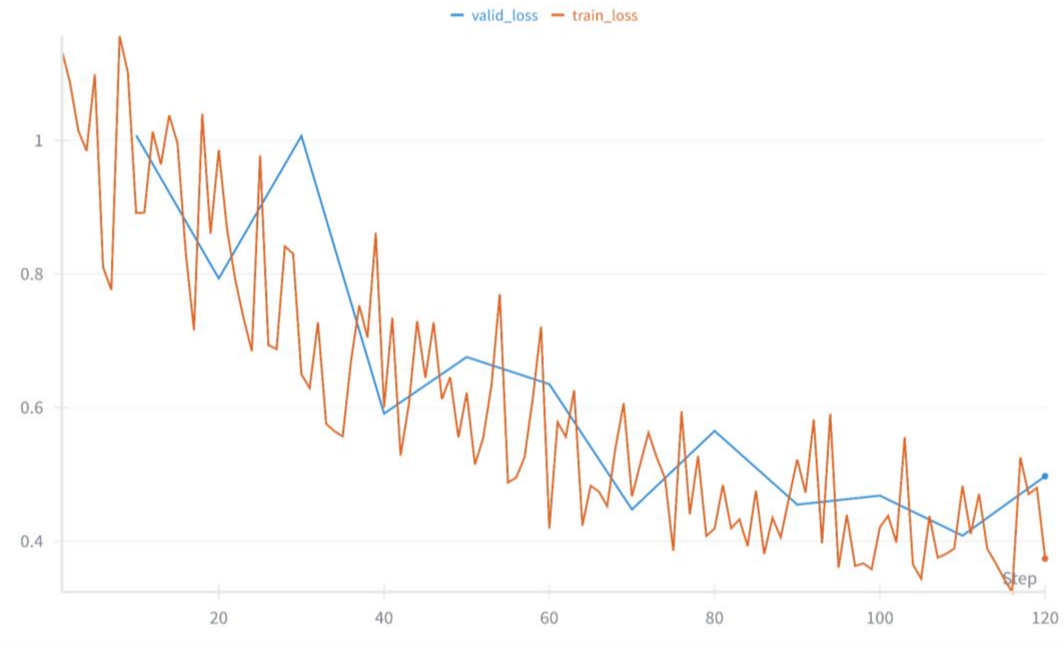


**Figure S1.** Training loss and validation loss. The X-axis is the training step, and the y-axis is the training and validation losses.

To understand the accuracy of the fine-tuned model, we used it to predict on the test set. The interaction of each instance was predicted three times. For a single comment, based on all predictions from the model, if there is an interaction detected, the instance will be classified "Yes". In reality, the proportion of eBird comments that contain bird interaction is much lower than that of irrelevant comments. Therefore, we tested the model five times on a heavily imbalanced dataset with only 67 instances labeled with "Yes" out of 2500 instances to further examine the model's capacity for detecting interaction.

The accuracy of interaction identification was described by two indicators: accuracy and the kappa coefficient. Accuracy measures the proportion of correctly predicted instances out of all instances. Kappa coefficient measures the agreement between the actual and predicted classifications, and is calculated as follows:

$$Kappa = \frac{Po - Pe}{1 - Pe} \text{ (S1)},$$

where $Po$ (Observed Agreement) represents the observed proportion of agreement, which is equal to accuracy. $Pe$ (Expected Agreement) represents the expected proportion of agreement by chance.

In the test set, the recall and accuracy for the interaction were 76.4% and 86.3%, respectively, as shown in Table S1. The value of 0.73 for the kappa coefficient indicates a high level of concordance between the forecast and actual results (0.6–0.8). Compared to the results of the test set, slightly higher recall (77.6%) and higher accuracy (97.9%) were obtained based on the imbalance dataset (Table 2). In addition, the lower kappa (0.66) still demonstrates a substantial agreement between prediction and observation. Therefore, the fine-tuned GPT-4o model is able to return stable detection of bird interaction with reliable accuracy.

**Table S1.** Confusion matrix for interaction identification with fine-tuned GPT-4o on the test set.

| **Interaction** | | **Ground truth** | | | |
|---|---|---|---|---|---|
| | | **Presence (Yes)** | **Absence (No)** | **Total** | **Precision** |
| Prediction | Presence (Yes) | 42 | 13 | 55 | 76.4% |
| | Absence (No) | 2 | 53 | 55 | 96.4% |
| | Total | 44 | 66 | 110 | |
| | Recall | 95.5% | 80.3% | | Overall: 86.3% |

Note: kappa: 0.73, indicating substantial agreement between prediction and observation

**Table S2.** Confusion matrix for interaction identification with fine-tuned GPT-4o on a larger dataset.

| **Interaction** | | **Ground truth** | | | |
|---|---|---|---|---|---|
| | | **Presence (Yes)** | **Absence (No)** | **Total** | **Precision** |
| Prediction | Presence (Yes) | 52 | 37 | 89 | 58.4% |
| | Absence (No) | 15 | 2396 | 2411 | 99.4% |
| | Total | 67 | 2433 | 2500 | |
| | Recall | 77.6% | 98.5% | | Overall: 97.9% |

Note: kappa: 0.66, indicating substantial agreement between prediction and observation

*Applying an LLM to categorize the interactions*

We first applied the fine-tuned GPT-4o model on all eBird comments in Michigan during the nonbreeding season (January, February, November, and December) in 2023 and 2024. Out of the 29,345 comments that contain interactions, we randomly selected a small subset of 485 comments for manual categorization. We then uploaded these comments to Claude Sonnet 4 using the following prompt, containing the same protocol used for manual categorization. The following protocol was adopted from the Avian Interaction Database (https://github.com/SpaCE-Lab-MSU/Avian-Interaction-Database). eBird comments are often short and contain vague descriptions of the specific interactions. Based on these characteristics, we simplified the number of interactions to 17 categories, added keywords that are indicators of certain interactions, and removed directionality (the asymmetric effect of certain interactions on the participants, e.g., predation will only benefit the predator) to assist with accuracy.

I need help classifying eBird comments by whether they describe interactions between bird species or not.

I am only interested in interactions between different bird species, so please exclude intraspecific interactions (interacting individuals of the same species, or a group of the same species flying or perching together) or interactions with other organisms that are not birds (mammals, insects, fish, amphibians, plants).

I define interactions as either direct engagement or co-occurrence. Below I define the specific types of interactions. The phrase before the colon is the type of interaction and should be used consistently in the output.

Aggression: aggressive behavior between species, but unsure of the motive.

Facilitation-mixed flocking: assign this if the comment indicates the species are deliberately associating, grouping, or flocking together. Signal phrases include: "with", "flocking with", "feeding together", "associating with", "moving together", "in a mixed flock", "together with". Example: "Continuing in area. Flew off southeast with Greater White-fronted Geese" -> Facilitation-mixed flocking (flying together signals intentionality)

Co-occur: assign this if the comment only shows the species are in the same area but there is no deliberate association. Signal phrases include: "near", "next to", "in the same tree as", "close to", "nearby". Example: "standing on pier with gulls" -> Co-occur (no intentionality; spatial proximity only)

Competition: aggressive behavior that is clearly from competition over space, territory, food, or any other resources.

Predation: predator eats prey, including unsuccessful attempts.

Nest predation: one species preying upon the young or eggs in a nest of another species.

Commensalism-call mimicry: when one species copies the song/call of another species.

Commensalism-scavenge: when one species consumes a previously deceased body of another species.

Commensalism-other: one species benefits and the other receives no effect, but the interaction does not fall into any other categories starting with commensalism above.

Kleptoparasitism: one species steals food or nest material from another species.

Brood parasitism: one species places its egg in another species' nest so the nesting species has to raise young of a different species.

Facilitation-comigration: when both species migrate together.

Facilitation-feeding: when both species feed together, either in mixed flocks or feeding aggregations.

Facilitation-communal nesting: when the species nest together with another species, usually during breeding season.

Facilitation-distress calls: when one species is under stress (e.g., presence of predator, or potential danger) and elicit a vocal response by another species.

Facilitation-other: when both species benefit from each other, but the interaction does not fall into any other categories starting with facilitation above.

Amensalism: one species causes harm to another species without any costs or benefits to itself, e.g., flushing another species from nest, roosting site, or foraging ground.

Mobbing: usually indicative of predation, where the smaller bird mobs the bigger bird. Can also happen when one species mobs another species to defend nest or young.

Given an input pair of the target bird species and the comment, identify the type of interaction.

Also, identify all of the bird species mentioned.

Please structure the output data in JSON format as shown below based on the given species and comment:

{{

"species1_common": string, # the same species listed in the "Species" field in the input. This is the focal species.

"species2_common": [list of strings], # the name(s) of the species that are involved in the interaction but are not the focal species as they appear in the comment (either full-name or abbreviation). If there are multiple, separate each with a comma.

"interaction": string # type of interaction, based on the definitions above. Only list the label with no additional text.

}}

--

Output Requirements:

- Return exactly one JSON object

- Each object must be valid JSON with double-quoted strings and keys.

- The "species2_common" field must always be a list — even if it contains only one species (e.g., ["Mallard"]).

- No trailing commas after the last item in the "species2_common" list.

- DO NOT include any explanatory text, markdown formatting, or backticks (`).

- Ensure the output is syntactically and structurally valid JSON that can be parsed by a JSON parser.

- Do not skip any entries, even if unsure — classify based on the best possible interpretation.

- Use the **full interaction label** as provided in the list above. Do not shorten or paraphrase it.

--

Example 1:

Species: Lapland Longspur

Comment: Mixed in with the larks, also giving classic twittering call. Will attach pics

Response:

{{

"species1_common": "Lapland Longspur",

"species2_common": ["larks"],

"interaction": "facilitation-mixed flocking"

}}

--

Example 2:

Species: Field Sparrow

Comment: Pink bill and legs, plain breast and gray face. Was in the large flock of ATSP and DEJU feeding in one of the fields.

Response:

{{

"species1_common": "Field Sparrow",

"species2_common": ["ATSP", "DEJU"],

"interaction": "facilitation-feeding"

}}

### Case study 2: Comments of a single pollinator from GBIF

The plant-pollinator case study consists of two steps: (1) evaluating the performance of LLM in categorizing plant-pollinator interactions with a smaller subset of comments, (2) applying the LLM to the entire dataset and comparing it with another data source (GloBI). Below, we describe step 1 in detail.

We manually categorized 400 comments for the butterfly *Vanessa annabella* on GBIF, then directly uploaded the comments to Claude Sonnet 4 using the following prompt containing the same protocol used in the manual categorization:

I need help classifying comments from observation of a single butterfly species by the type of biotic interactions it engages in. If the comments are not in English, translate them into English first.

I am only interested in interactions between the butterfly and other plants. I define interactions as either direct engagement or co-occurrence. Below I define the specific types of interactions. The phrase before the colon is the type of interaction and should be used consistently in the output.

Pollination: direct evidence that the adult butterfly is visiting a flower of a plant species. Key phrases include: nectar/nectaring, visiting the flower, feed/feeding/forage/foraging on flower, working on, using flower, stopping on flower, attracted by flower.

Herbivory: direct evidence that the larvae of the butterfly (i.e., caterpillars) are feeding on a host plant species, or eggs of the butterfly are found on the host plant species, or the adult butterfly is laying eggs on the host plant species.

Co-occur: when the butterfly is associating with another species but does not fall into any of the categories above.

No interaction: when the comment does not contain any information of the butterfly engaging with other plant species. Examples include simple description of location, weather, appearance of the butterfly, or butterflies engaging with each other.

Output each result in JSONL format, one JSON object per line, with the fields: "id" (original record ID), "species2" (the plant species name(s) mentioned in the comment, written exactly as they appear; if multiple, separate with commas; if none, leave as an empty string) and "interaction" (one of "Pollination", "Herbivory", "Co-occur", or "No interaction").

Example: {"id": 923916653, "species2": "milkweed", "interaction": "pollination"}

We used a strict definition of pollination that the comment must explicitly mention the flower-visiting behavior. For example, “nectaring on lantana” would be considered pollination, while “on lantana” would only be considered as co-occurrence. Although this strict definition will likely categorize many actual pollination interactions as co-occurrence, it ensures the quality of the output data. Contrary to case study 1, we did not fine-tune the LLM because Claude Sonnet 4 does not allow for fine-tuning, and smaller, open-source models did not achieve the same level of performance as Claude Sonnet 4 even after fine-tuning on the training dataset.
